# Metabolic adaptations underlying genome flexibility in prokaryotes

**DOI:** 10.1101/415182

**Authors:** Akshit Goyal

## Abstract

Even across genomes of the same species, prokaryotes exhibit remarkable flexibility in gene content. We do not know whether this flexible or “accessory” content is mostly neutral or adaptive, largely due to the lack of explicit analyses of accessory gene function. Here, across 96 diverse prokaryotic species, I show that a considerable fraction (~40%) of accessory genomes harbours beneficial metabolic functions. These functions take two forms: (1) they significantly expand the biosynthetic potential of individual strains, and (2) they help reduce strain-specific metabolic auxotrophies via intra-species metabolic exchanges. I find that the potential of both these functions increases with increasing genome flexibility. Together, these results are consistent with a significant adaptive role for prokaryotic pangenomes.

**Author Summary:** Recent and rapid advancements in genome sequencing technologies have revealed key insights into the world of bacteria and archaea. One puzzling aspect uncovered by these studies is the following: genomes of the same species can often look very different. Specifically, some “core” genes are maintained across all intraspecies genomes, but many “accessory” genes differ between strains. A major ongoing debate thus asks: do most of these accessory genes provide a benefit to different strains, and if so, in what form? In this study, I suggest that the answer is “yes, through metabolic interactions”. I show that many accessory genes provide significant metabolic advantages to different strains in different conditions. I achieve this by explicitly conducting a large-scale systematic analysis of 1,339 genomes across 96 diverse species of bacteria and archaea. A surprising prediction of this study that in many ecological niches, co-occurring strains of the same species may help each other survive by exchanging metabolites exclusively produced by these different accessory genes. More pronounced gene differences lead to more underlying metabolic advantages.

## Introduction

Prokaryotes exhibit remarkable genome flexibility, with strains from the same species often containing dramatically different gene content^1-4^. Intraspecific differences in gene content are often characterized by a “core” genome (genes common to all strains) and “accessory” genome (genes found in a fraction of strains)^5^. While the core genome might represent a set of species-specific indispensable genes, we do not yet understand whether the accessory genome of a species is the result of neutral or adaptive evolution. Indeed, this is the subject of an ongoing debate: do the majority of prokaryotic accessory genes have negligible or positive fitness effects, i.e. are they neutral or adaptive?

Recent population genetics arguments support roles for both neutral and adaptive evolution as possible factors driving accessory genome evolution^6-9^. For example, microbial species with more accessory genes also tend to have larger effective population sizes, as expected of genetic variation in a population under neutral evolution^9^. On the other hand, models in which microbial genomes evolve in large, migrating populations, suggest that acquired genes can often be beneficial, as expected under adaptive evolution^7^. However, these studies have only addressed broad aspects of microbial populations such as effective population size, migration, and the fitness effects of gene loss and gain. In response, subsequent criticisms of these studies have strongly expressed the need for more functional, gene-explicit and ecological analyses^10-11^. Here I present the first such systematic analysis of 96 phylogenetically diverse prokaryotic species, which suggests that prokaryotic accessory genomes often provide significant metabolic benefits.

I chose to study metabolism as a possible explanatory factor for three reasons. (1) Metabolic genes dominate the functional content of accessory genomes^12^ (supplementary figure 8a). (2) Metabolic interactions between microbes—especially interdependencies—can often be adaptive^13-14^. For instance, microbes that obligately cross-feed, i.e. that critically depend on exchanging metabolites with each other, can grow faster than their wild-type counterparts^13^. Such a fitness benefit can also drive genomes, in many cases, to lose genes and become metabolically dependent^14,39^. If different genes are lost between different conspecific strains, this can lead to both metabolic interdependence, as well as accessory genomes (since different strains will have different metabolic repertoires)^15^. (3) Databases such as KEGG contain already-curated genomes for several fully-sequenced strains. KEGG contains high-quality gene and reaction annotations, allowing us to accurately predict the biosynthetic capabilities of each strain under different conditions^16^.

In this study, I ask to what degree accessory genes can metabolically benefit conspecific strains. For this, I have used genome-scale metabolic network reconstructions of 1,339 prokaryotic strains (corresponding to 92 bacterial and 4 archaeal species) from the KEGG database over 59 distinct nutrient environments. In general, my analyses reveal two beneficial roles for the accessory metabolic content of prokaryotes. First, I find that the accessory genome of most species harbours extensive biosynthetic potential, with several accessory genes providing strains with additional nutrient utilization abilities. Second, I find that pairs of strains from the same species often display a remarkable potential for metabolic interdependence, which scales with the amount of accessory genome content. These interdependencies have the ability to alleviate strain-specific auxotrophies in a particular niche through the exchange of secreted metabolites. My results are, from a metabolic standpoint, consistent with a possible adaptive evolution of accessory genomes.

## Results and Discussion

To obtain a large set of species pangenomes, I first collected a list of all prokaryotic species in the KEGG GENOME database, and filtered those that had complete genomes for 5 or more conspecific strains. This gave me 1,339 genomes (96 species), which I used in all my subsequent analyses (supplementary table 1). To account for potential biases due to uneven phylogenetic sampling, I verified that a more restrictive choice of one species per genus did not significantly impact my results (55 species; supplementary figure 1). For each strain, I then extracted all annotated genes and metabolic reactions from the KEGG GENES and REACTION databases, respectively. To quantify accessory genome content for every species, I used the well-studied genome fluidity measure, *φ*^17^. For this, I calculated, across each pair of conspecific strains, the fraction of all genes in the pair that were unique to each strain. The average of this fraction over each species gave me its genome fluidity *φ* (see Methods).

I constructed metabolic networks for every individual strain, where each network contained the set of reactions corresponding to the strain’s genome in KEGG. I included gap-filled reactions when curated models were available^18^, though I verified that their addition did not impact my results (supplementary figure 2). I used these reaction networks to infer the biosynthetic capabilities of each strain under several different conditions. To define these conditions, I selected 59 different carbon sources, previously shown and commonly used to sustain the growth of diverse microbial metabolisms in laboratory experiments^19-21^ (supplementary table 2). I associated with each carbon source a different nutrient environment or condition. In each condition, I included exactly one of the 59 carbon sources, say glucose, along with a set of 30 commonly available metabolites, which I assumed were always available (for example, water and ATP; supplementary table 3).

To assess biosynthetic capability, I curated a list of 137 crucial biomass precursor molecules, often essential for growth (hereafter, “precursors”) from 70 experimentally verified high-quality metabolic models^22^ (supplementary table 4). Finally, to calculate what each strain could synthesize in a particular environment, I used a popular network expansion algorithm: called scope expansion^23-24^. This algorithm determines which metabolites each strain can produce — its “scope” — given an initial seed set of already available metabolites. To start with, only those reactions whose substrates are available in the environment can be performed, and their products constitute the initial set of metabolites that can be produced. These metabolites can then be used as substrates for new reactions that can then be performed, and step by step, more metabolites can be produced. When no new reactions can be performed, the algorithm stops, giving the full set of metabolites that could be synthesized in the given environment. Such a calculation sidesteps the need for arbitrary assumptions of binary (yes/no) growth and optimality typically used in more complex metabolic modeling approaches such as flux balance analysis^25^ and is well-known for its ability to infer what metabolic networks can synthesize in diverse conditions^26-27^.

I first investigated the capabilities of individual strains. Specifically, I was interested in the extent to which the accessory genes in each strain expanded the set of precursors that could be synthesized. For each species, I calculated, via network expansion, the list of precursors that could be produced per strain per condition. I then counted how many unique precursors each strain could synthesize across all conditions, i.e. by the accessory genes alone. From this, I computed, for every species, its accessory metabolic capacity *α*, defined as the average number of precursors (per strain per condition) produced exclusively due to the accessory genome.

I found that while for 18 species this quantity was zero, for the majority of prokaryotic species (81%), this number lay between 0.1 and 15.0 (mean 3.1; median 2.0). Further, *α* scaled positively with genome fluidity *φ* (Spearman’s rho = 0.44; *P* value = 7 x 10^-6^; figure 1).

**Figure 1.**
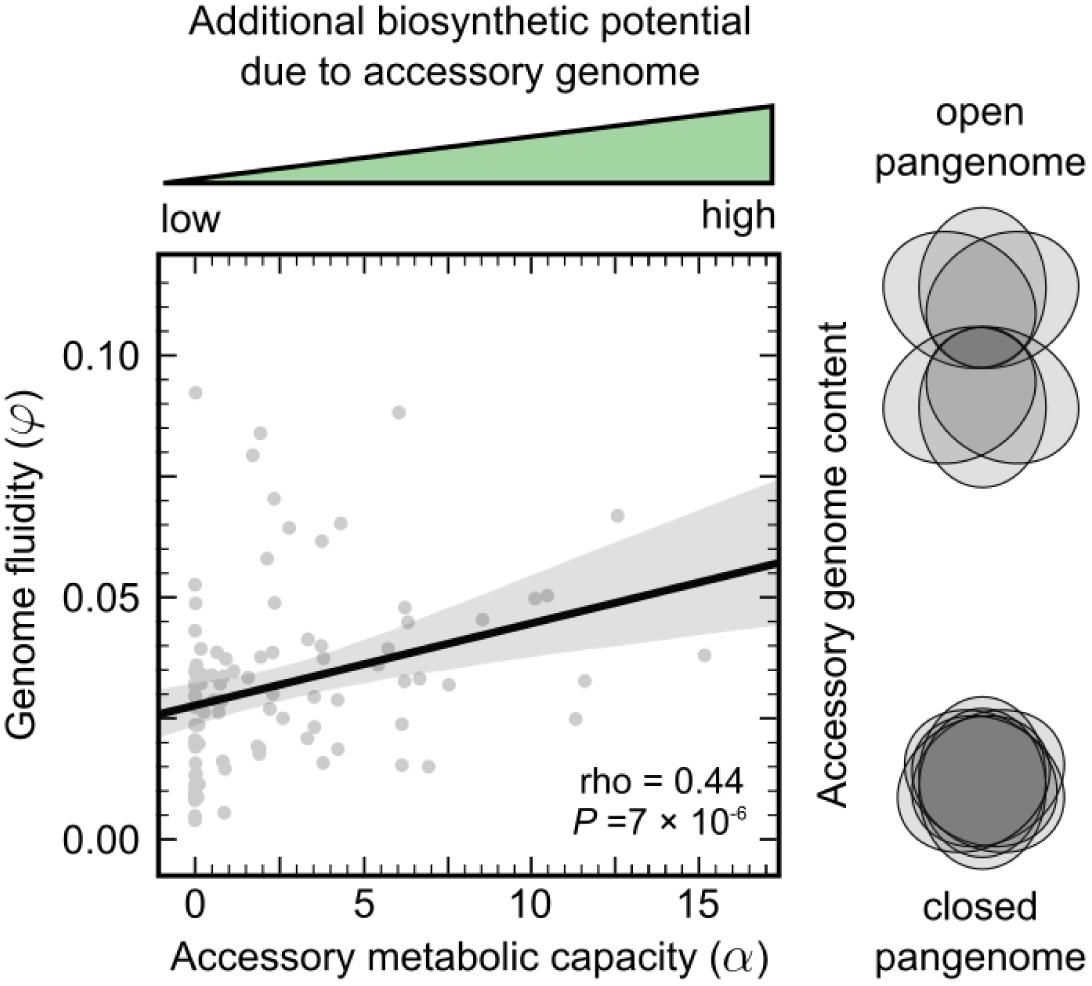
The accessory genomes of prokaryotes harbour extensive biosynthetic potential. Scatter plot of genome fluidity *φ* versus accessory metabolic capacity *α* for the 96 prokaryotic species in this study. Each point represents the average number of precursors that could be synthesized by the accessory genome content alone in each strain of a species. The Venn diagrams on the right provide a schematic representation of open pangenomes (high *φ*, small core) versus closed pangenomes (low *φ*, large core). The solid black line represents a linear regression and the gray envelope around it, the 95% prediction interval. rho corresponds to Spearman’s nonparametric correlation coefficient and the *P* value to a one-way asymptotic permutation test for positive correlation.

Since I observed that the accessory genome of different strains typically imparted different biosynthetic capabilities to different strains, I wondered if, in the same conditions, metabolic interactions between conspecific strains could further expand these capabilities. This could, for instance, indicate a potential dependence of an auxotrophic strain on another strain, i.e. a strain that cannot produce a crucial precursor in an environment. Such auxotrophies have been previously shown for example, in different strains of *Escherichia coli* co-inhabiting the human gut^28^.

For each pair of conspecific strains in each condition, I calculated a metabolic dependency potential (MDP), defined as the average number of new precursors each strain has the potential to synthesize when grown as a pair versus alone. Here I assessed, in every pair, which metabolites that could be produced and secreted by one strain could subsequently allow the production of a new precursor in the other strain that it would not otherwise be able to make (i.e. was auxotrophic for). Note that this method does not count those metabolic interactions that can provide extra (functionally redundant) pathways to produce a precursor and supplement growth, and is thus more likely to represent actual or obligate dependencies. I verified that my approach can successfully predict such obligate dependencies by comparing with some well-documented intra- and inter-species pairs^13-14,29-31^ (see Methods) (supplementary figure 3).

I found that while I could not detect any dependency potential for 17 species, surprisingly, the majority of species (82%) showed an MDP per strain per condition between 0.1 (for *Bacillus thuringiensis*) and 3.3 (for *Ralstonia solanacearum*), with a mean 1.7 and median 1.4. Interestingly, the 17 species for which I could not detect any MDP matched those with zero *α* (the leftover *Legionella pneumophila* showed low MDP = 0.5). Over all tested pairs with detected dependency potential (48%), commensal interactions were more common than mutualisms (29% versus 19%; figure 2b). This is because, in species with detected dependencies, not all pairs show dependency potential (on average 46% conspecific pairs do). The auxotrophies relieved by these dependencies varied from those for amino acids, vitamins, carbohydrates, and organic acids, among others (figure 2c).

**Figure 2.**
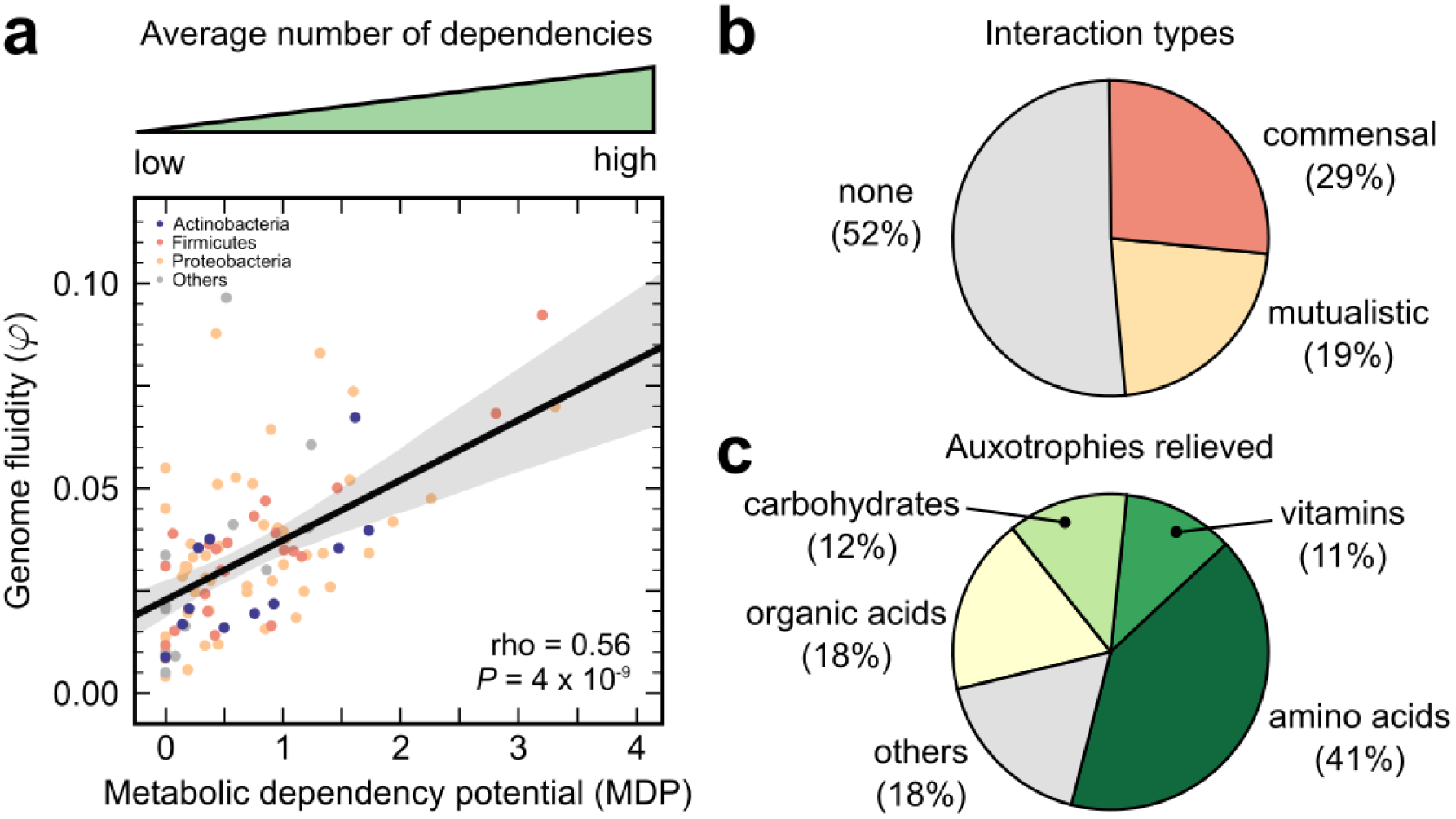
Conspecific metabolic dependencies scale with accessory genome content. **a,** Scatter plot of genome fluidity *φ* versus conspecific metabolic dependency potential (MDP) for the 96 prokaryotic species in this study. Each point represents the average number of dependencies detected per strain per condition across all conspecific pairs for one species. Colours represent each species’ phylum-level taxonomic identity. The solid black line represents a linear regression and the gray envelope around it, the 95% prediction interval. rho corresponds to Spearman’s nonparametric correlation coefficient and the *P* value to a one-way asymptotic permutation test for positive correlation. **b,** Pie chart of the types of interactions detected in each conspecific pair. **c,** Pie chart of the types of auxotrophies that the detected dependencies relieve due to pairwise growth. Each of the 137 precursors belongs to a unique chosen category (supplementary table 4).

Strikingly, like *α*, MDP also scaled positively with genome fluidity *φ*, suggesting that greater amounts of accessory content can potentially sustain more conspecific metabolic dependencies (Spearman’s rho = 0.56, *P* value = 4 x 10^-9^; figure 2a). For both *α* and MDP, considering medians instead of means did not impact my results (for *α*: Spearman’s rho = 0.38, *P* value = 10^-4^; for MDP: Spearman’s rho = 0.51, *P* value = 10^-7^; supplementary figures 4 and 5). To verify that such potential conspecific dependencies are indeed ecologically realizable, I repeated my analysis, this time restricting it to those genomes, which were known to co-occur in microbial communities (29 strains across 14 species; see Methods). I found that my observed trend was still valid, namely MDP still scaled with *φ*, suggesting that several auxotrophies may indeed be reduced through within-species metabolic exchanges in nature (Spearman’s rho = 0.56, *P* value = 0.03; supplementary figure 6).

Given the extent to which I detected the potential for obligate metabolic interactions between conspecific strains, I wondered whether such interactions are possibly common among prokaryotes. For this, I extended my study to analyze metabolic dependency potential between inter-specific strains (see Methods). I found that, indeed, strains from all species can metabolically depend on strains from at least one other species to alleviate potential auxotrophies across many different environments (with MDP ranging from 1.6 to 3.2, with mean 2.2; supplementary figure 7). Interestingly, I found that these interspecific metabolic interactions often involve accessory metabolic genes as well.

Taken together, my results suggest that a considerable fraction of prokaryotic accessory genomes contains potentially beneficial metabolic functions (upto 70% of accessory genes per strain across my study, with median 40%; see supplementary figure 8d). Specifically, I found that the accessory “metabolome”: (1) expands a genome’s biosynthetic potential, possibly allowing for niche-specific adaptations^28, 32-33^; and (2) reduces potential auxotrophies via obligate metabolic interactions, also explaining how conspecific strains can coexist despite high competitive potential34-37. The accessory genes that impart these functions are often different (median overlap 10%), suggesting that these are indeed distinct, non-redundant benefits. Moreover, apart from these additional biosynthetic abilities, the y-intercept for genome fluidity (at *φ* = 0.03 for both *α* and MDP) provides an estimate of metabolic redundancies (such as extra pathways).

My findings may additionally help explain the following observations: (1) metabolic functions are enriched in accessory genomes (median 50% in accessory versus 38% in core; supplementary figure 8c); and (2) the variation in accessory metabolic genes exceeds the variation in genes of many other functions (metabolic variation being dominant in 81% of examined cases; supplementary figure 8b).

Previous studies have suggested that the evolution of metabolic dependencies likely occurs via adaptive gene loss^39-40^ (e.g. the Black Queen hypothesis). Such a mechanism suggests that metabolic dependency evolution can often lead to reduced genome sizes, but makes no comment on genome flexibility (i.e. gene content variability). My results also indicate that metabolic dependency evolution can impact genome flexibility as well. Specifically, more flexible genomes (with more variable gene content) are more likely to display a potential for metabolic interactions.

Can stochastic accessory gene turnover explain these results? To test this, I repeated my study with randomly assembled pangenomes. Within each species, I retained the core genes in every strain and shuffled the accessory genes between strains (see Methods). During this randomization, I preserved the observed within-species genome size distribution, strain number distribution, and ensured that any change in species’ genome fluidity was insignificant. I found that not only did this significantly diminish the metabolic benefits observed in each species, both measurements of *α* and MDP yielded non-significant correlations, suggesting that the mere presence of additional accessory genes is unlikely to explain my observed trends (for *α*: Spearman’s rho = 0.01, *P* value = 0.9; for MDP: Spearman’s rho = 0.17, *P* value = 0.1; supplementary figures 9a and 10a). The measured benefits remained lower than observed, even when I shuffled known operons of genes together instead of shuffling genes one by one (supplementary figures 9b and 10b; see Methods). I believe this is because often, prokaryotic operons do not contain complete metabolic pathways, but instead parts of them (in my data set, each metabolic operon encoded 1.5 reactions on average, while pathways typically had 4 to 5 steps). Collectively, this suggests that accessory gene acquisition is consistent with the coordinated gain of functional and beneficial pathways, which I believe provides further support for the accessory genes being maintained for adaptive reasons.

To summarize, here I addressed the debate on whether the accessory genomes of prokaryotes are beneficial. I found that, indeed, large fractions (about 40%) of the prokaryotic accessory gene pool can contribute to metabolic benefits. Specifically, such genes can allow microbes to produce a larger repertoire of crucial molecules, and facilitate the exchange of important metabolites. Since these functions can improve growth in many habitats, my results suggest that maintaining accessory genes can be adaptive. Note, however that my analyses are only capable of detecting obligate metabolic dependencies and biosynthetic potential, and do not consider signaling, regulation, metabolic redundancy, etc. that could also play important functional roles and might indicate potential benefits due to additional accessory genes. Further work might also explain accessory genomes in those species, where I could not detect additional metabolic functions, if such roles are indeed there. Moreover, even in the context of metabolism, more detailed metabolic models, when available, may be used to probe even more precise fitness effects of intraspecies metabolic variability, including the effect of higher-order interactions. However, these studies would require knowledge of a large number of parameters such as reaction kinetics, thermodynamics, and exact strain biomass compositions before they are feasible. Finally, systematic measurements of the fitness effects of all accessory genes, metabolic and otherwise, are needed for more complete estimates of the fraction of accessory genomes consistent with adaptive versus neutral evolution.

## Methods

### Acquiring pangenomes and metabolic networks from KEGG

I used the KEGG GENOMES database^16^ to extract a list of all prokaryotic species with complete genomes for 5 or more strains. This yielded a list of 1,339 strains or genomes corresponding to 96 species (92 bacteria, 4 archaea), which I used for all subsequent analyses (see supplementary table 1 for the full list of species and strains, along with their taxonomic classification). For each strain with a unique genome abbreviation, I extracted the full set of annotated genes under the KEGG GENES database and reactions under the KEGG REACTION database using an in-house Python script. I also extracted the full list of reactions with their stoichiometries and participating metabolites in the database. The metabolic reaction network for each strain was considered to be the complete set of annotated reactions detected in that strain’s genome in KEGG. Note that my analyses systematically ignore genes without known functions.

### Adding gap-filled reactions from Model SEED

For strains for which genome-scale metabolic reconstructions were available in the Model SEED database^18^, I also included gap-filled reactions. Specifically, I extracted the list of all gap-filled reactions for 130 genomes from table S3 in ref. 18. I mapped all genomes from this table to KEGG genomes by matching strain names, and all reaction IDs to KEGG reactions by searching the Model SEED database online (https://modelseed.org/biochem/reactions) using a custom Python script. This resulted in a total of 562 gap-filled reactions, spread across 22 genomes (20 out of 96 species; supplementary table 6). I then added these reactions to the metabolic networks already constructed via KEGG. Separately, I verified that adding these gap-filled reactions did not impact my results (supplementary figure 2).

### Defining nutrient environments or conditions

For nutrient environments or conditions, I selected a set of 59 diverse carbon sources known to sustain microbial biomass and energy synthesis from previous genome-scale metabolic studies of phylogenetically broad species^19-21^ (supplementary table 2). Every condition was assumed to contain one of these carbon sources (such as glucose and maltose), along with a set of 30 commonly available metabolites (assumed to be present in all conditions, such as water, oxygen and ATP), similar to the aforementioned studies (supplementary table 3). To infer biosynthetic potential, I separately collected a set of all prokaryotic species biomass compositions and their constituent metabolites from high-quality experimentally verified metabolic models in the BiGG database22. I curated from this a list a union of 137 biomass precursors across diverse microbial metabolisms (supplementary table 4).

### Network expansion algorithm

To infer what each strain could synthesize in each nutrient environment or condition, I used a well-documented network expansion algorithm — scope expansion^23-24^. Briefly, this algorithm is given a reaction network (one from each genome) and an initial “seed” set of available metabolites (each nutrient environment). It first determines which reactions can be performed by the network using only the nutrients in the environment. I assume that metabolites that are products in this initial set of reactions can be synthesized by the network, and can be subsequently used as reactants in new reactions. Again, I consider that the products of such new reactions can be synthesized by the network, and may allow additional new reactions to be performed. This continues step by step, till no no new reactions can be performed. All metabolites that can be produced over all such steps are defined as the “scope” of the metabolic network, i.e. I assume that these metabolites can be synthesized by the reaction network from the initial nutrients in the environment.

### Calculating genome fluidity

I calculated genome fluidity *φ* as prescribed in a previous study^17^, using a custom Python script. For every genome, I considered each constituent gene’s KO number as its unique identifier. Then, to estimate *φ* for every species, I calculated, for all conspecific pairs, the ratio of the number of genes unique to a strain in the pair to the total number of genes in their sum. The average over all pairs for a species was considered its genome fluidity *φ*. Note that though using KEGG orthologous groups underestimates the exact values of *φ*, my estimates still scale well with previously reported values9 (Spearman’s rho = 0.60, *P* value = 7 x 10^-5^; supplementary figure 11).

### Calculating accessory metabolic capacity

I calculated an accessory metabolic capacity *α* for every species. For each conspecific strain, I first calculated, using the network expansion algorithm described, the scope of each of the 1,339 reaction networks across all 59 conditions. Then, species by species, for every condition, I calculated a “core” metabolome, i.e. metabolites that were present in the scope of every conspecific strain. I then explicitly removed these metabolites within every species from the scope of each strain and counted how many precursors remained in the corresponding “accessory” metabolome of every strain across all conditions. This gave me a number of additional precursors that could be synthesized per strain per condition for each species, and was defined as the species’ accessory metabolic capacity *α* (supplementary table 5).

### Calculating metabolic dependency potential between conspecific pairs

I calculated a metabolic dependency potential (MDP) for every species. For this, I considered within each species, all conspecific pairs across all 59 conditions. For each pair, I calculated the scope for both strains first in “monoculture”, i.e. when grown alone. I then calculated, for every metabolite that could be produced by one strain but not the partner strain, whether or not its secretion could alleviate an auxotrophy in the partner. I specifically considered auxotrophies only for the 137 key precursors I had selected. I then counted each alleviated auxotrophy as a potential metabolic dependency, and the average number of dependencies (per strain per pair per condition) for each species was defined as its metabolic dependency potential, or MDP (supplementary table 5).

### Calculating metabolic dependency potential between inter-specific pairs

To quantify the extent of metabolic interactions between inter-specific strains, I calculated a separate metabolic dependency potential for every species. For each species, I paired each conspecific strain with 25 randomly chosen strains from other species, also picked at random. For all inter-specific pairs generated this way, I calculated metabolic dependency potential using the same method as described above, for conspecific pairs. In this way, the average number of dependencies identified per strain per condition was defined as the inter-specific metabolic dependency potential, or MDP (supplementary table 5).

### Determining strain co-occurrence from microbial community data

To test whether the conspecific metabolic interactions detected in my MDP analysis could be realized in natural microbial communities, I analyzed genome co-occurrence data from Chaffron et al38. These data list all 16S rRNA sequences co-detected across several microbial community samples. Here, sets of sequences are clustered into operational taxonomic units (OTUs) corresponding to different sequence similarity thresholds. To map these OTUs to the genomes in my study, I first obtained 16S rRNA sequences for all the 1,339 genomes I analyzed from KEGG. When multiple sequences were available for a given genome, I used the longest sequence and maped that as the unique 16S identifier for that strain. Then, using BLAST, I mapped OTUs in the co-occurrence data to the genomes in my study (where OTUs were binned with a sequence similarity threshold of 99%). Here, I used the BLAST bit score as my assignment criterion. I used the 689 genomes that could be mapped this way for further analysis. Here, across all microbial community samples, I asked which conspecific genomes co-occurred in at least one sample — from which I found 29 genomes corresponding to 14 species (supplementary table 7). I then repeated my metabolic dependency potential analysis for these conspecific strains, as described above.

### Testing for the role of stochastic accessory gene turnover

To test if my observed correlations between *φ* and *α* as well as *φ* and MDP could be explained by random accessory gene turnover, I repeated my study with a “randomly assembled” pangenome dataset. I randomized genomes species by species. I first collected all available genomes for a species, and picked a random pair of these. I then shuffled the accessory genes in this pair in two ways: (1) gene by gene, and (2) operon by operon.

When shuffling gene by gene, for each strain pair, I randomly picked two genes, one from each strain in the pair, and swapped them. I repeated this several times before picking another conspecific pair from the same species. The number of swaps per pair was chosen such that each accessory gene was swapped once on average. I verified that the exact number of swaps does not affect my results. By the end of this process, I had a new set of genomes which had undergone “stochastic accessory gene turnover”. Note that in order to avoid any potential biases, this process preserves the observed genome sizes and strain numbers while only slightly affecting genome fluidity. I then repeated my *α* and MDP calculations for these “shuffled” genomes. This would test if the mere acquisition of extra genes from a species’ accessory genome could allow the expanded biosynthetic potential and metabolic dependencies observed in the data.

To identify operons, I used the ProOpDB database, which lists operon compositions for more than 1200 prokaryotic genomes^41^. I found that this database had operon compositions available for 795 strains across 64 of the species in my study, which I used for the operon shuffling analysis (supplementary table 8). Here, when shuffling operons, I used a similar method as when shuffling genes, but instead of swapping merely randomly chosen genes from a pair of strains, I identified which operon they belonged to in their respective strain’s genome, and swapped all genes in those operons across the pair. I repeated these operon swaps several times for each strain pair, and for several pairs, at the end of which, I had another new set of randomly shuffled genomes.

### Comparing predicted dependencies with experimentally verified pairs

To test if my metabolic dependency potential (MDP) measure could accurately predict metabolic dependencies between different pairs, I compared its performance on genome-scale metabolic networks corresponding to some well-studied experimentally verified metabolically dependent pairs. Specifically, I considered 2 conspecific and 4 inter-specific pairs. For conspecific dependencies, I used 2 *Escherichia coli* cross-feeding pairs^13^ and for inter-species, I used (1) a *Desulfovibrio vulgaris* and *Methanococcus maripaludis* pair^29^; (2) an *E. coli* and *Acinetobacter baylyi* pair14; (3) an *Lactobacillus bulgaris* and *Streptococcus thermophilus* pair^30^; and (4) a *Bifidobacterium longum* and *Eubacterium rectale* pair^31^. In all cases, I obtained the metabolic models for the closest available strains from KEGG and, when needed, modified the genes present to best match those described in the respective studies. I then used my MDP approach as described to infer which potential dependencies were detected in each pair for the specific conditions mentioned in each study. I found that my method could accurately identify the extent of dependencies (number of auxotrophies) and interaction directionality (commensal or mutualistic interactions; supplementary figure 3).

### Statistics

To calculate correlation coefficients throughout the study, I used Spearman’s nonparametric rho, and for *P* values, I used a one-way asymptotic permutation test for positive correlation. All statistical tests were performed using standard implementations in the SciPy (version 0.18.1) and NumPy (version 1.13.1) libraries in the Python programming language (version 3.5.2). Linear regression and prediction interval calculations were performed using the Seaborn library function regplot (version 0.7.1).

### Data and code availability

All computer code and extracted data files used in this study are available at the following URL: https://github.com/eltanin4/pangenomedep.

## Acknowledgements

This work was supported by the Simons Foundation. I thank Sergei Maslov, Saurabh Mahajan, Rohini Subrahmanyam, Vishaka Datta, Sandeep Krishna, Deepa Agashe, and Sankarshan Talluri for discussions and comments on the manuscript. I am especially grateful to Christian Kost and two anonymous reviewers for their suggestions and comments on earlier drafts of this manuscript.

## Supplementary Figures

**Supplementary Figure 1.**
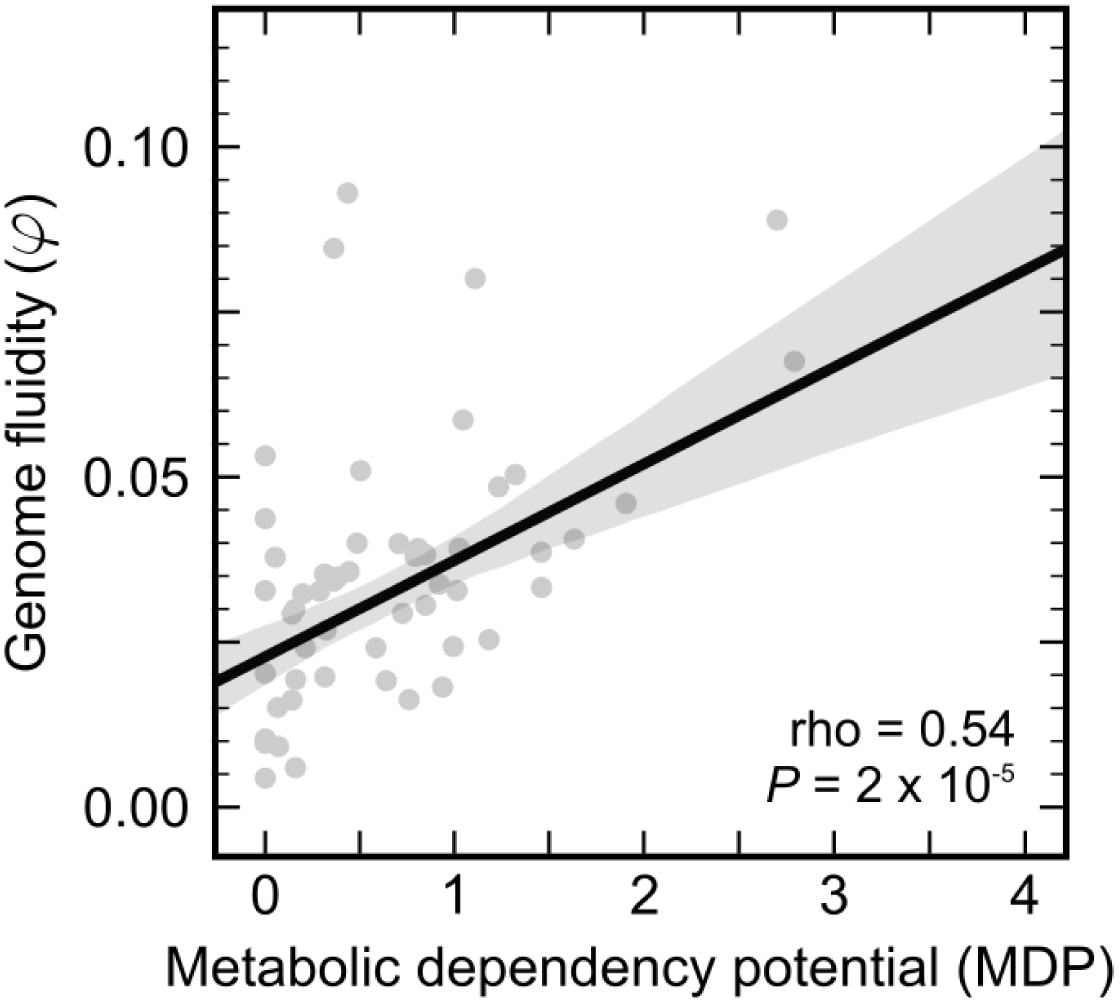
Phylogenetic bias in species set does not impact my key result. Scatter plot of genome fluidity *φ* versus conspecific metabolic dependency potential (MDP), similar to figure 2, but to minimize phylogenetic bias, here I only included one species per genus. This resulted in 55 species (51 bacteria, 4 archaea). Each point represents the average number of dependencies detected per strain per condition across all conspecific pairs for one species. The solid black line represents a linear regression and the gray envelope around it, the 95% prediction interval. MDP still increases significantly with increasing genome fluidity.

**Supplementary Figure 2.**
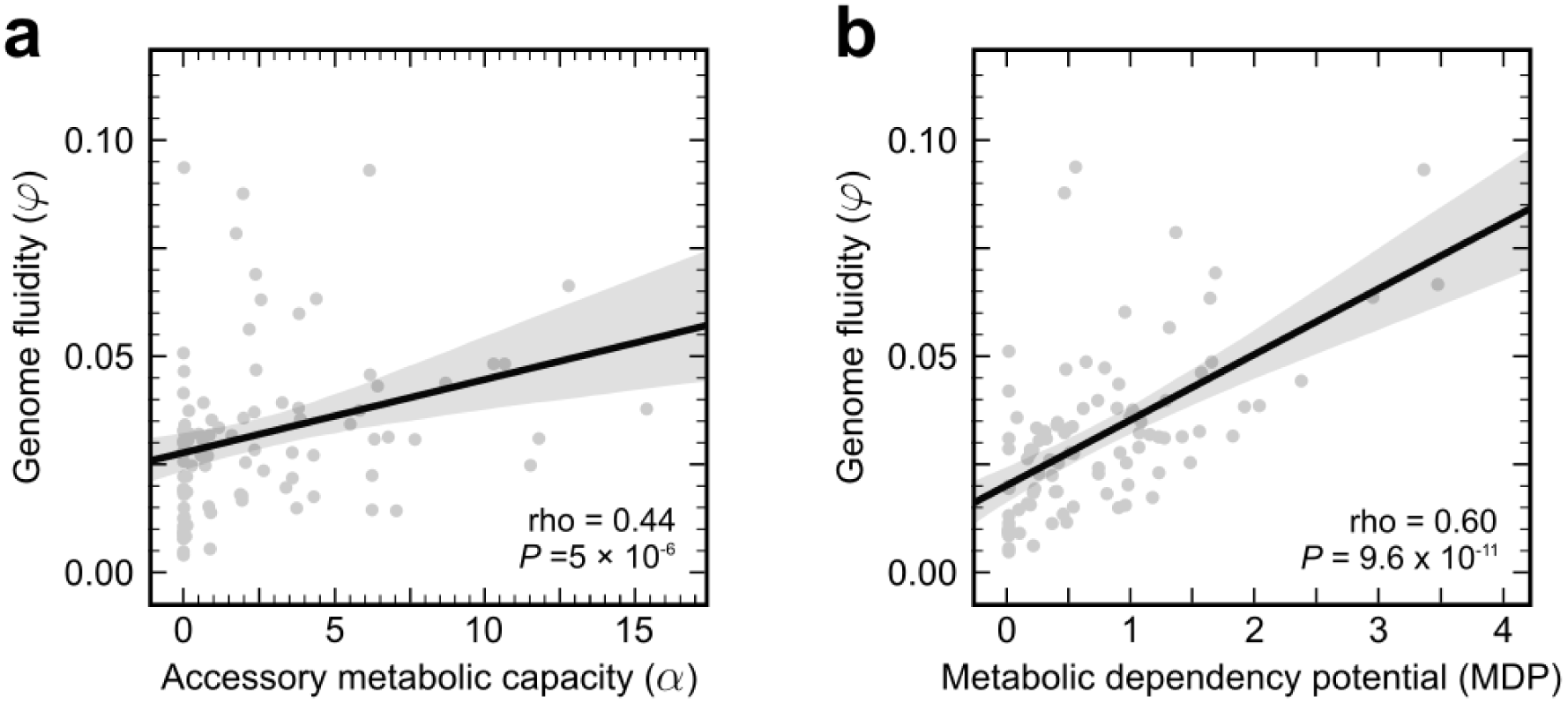
Using gap-filled metabolic models does not impact my results. **a,** Scatter plot of genome fluidity *φ* versus accessory metabolic capacity *α* for the 96 prokaryotic species in this study, similar to figure 1; and **b,** scatter plot of genome fluidity *φ* versus conspecific metabolic dependency potential (MDP), similar to figure 2; except without the addition of any gap-filled reactions (see Methods). This eliminates any potential bias that may arise from adding gap-filled reactions. Each point represents one species, both solid black lines represent linear regression, and the gray envelopes around them, 95% prediction intervals. Both observed trends are qualitatively unaffected.

**Supplementary Figure 3.**
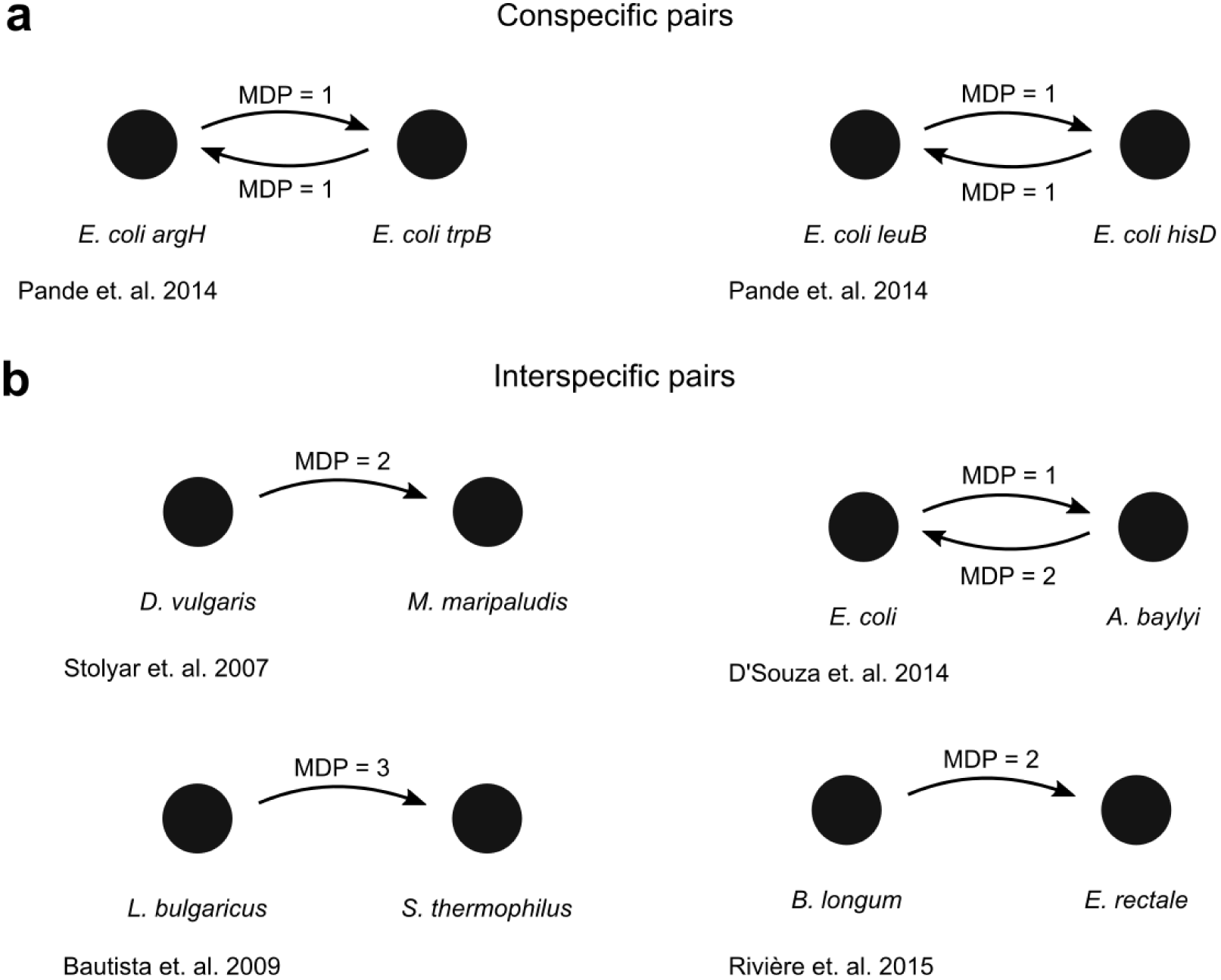
Metabolic dependency potential (MDP) accurately captures experimentally verified dependencies. For **a,** 2 conspecific and **b,** 4 interspecific pairs of prokaryotes, I verified that my approach to infer and measure metabolic dependencies using KEGG-annotated metabolic reaction networks (see Methods) can predict both the number of dependencies between each microbe, as well as the correct interaction type (commensalism, as in the top-left pair in b; and mutualism, as in the top-right pair).

**Supplementary Figure 4.**
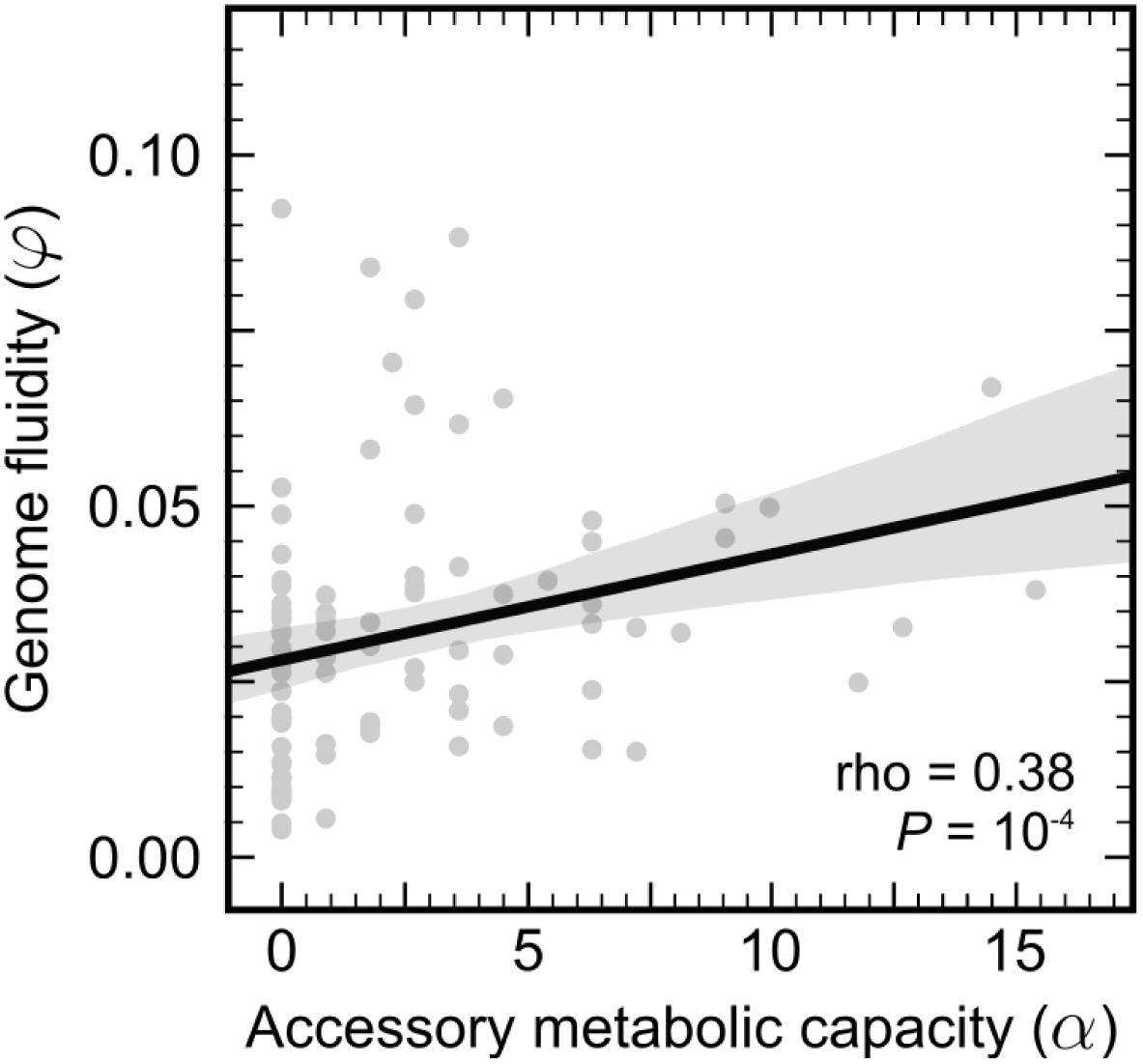
The impact of considering medians instead of means for *α*. Same as figure 1, except here to calculate *α*, instead of using the mean number of precursors produced by each individual strain’s accessory genome, I considered medians. The solid black line represents a linear regression and the gray envelope around it, the 95% prediction interval. rho corresponds to Spearman’s nonparametric correlation coefficient and the *P* value to a one-way asymptotic permutation test for positive correlation. This choice does not significantly impact my results for *α*.

**Supplementary Figure 5.**
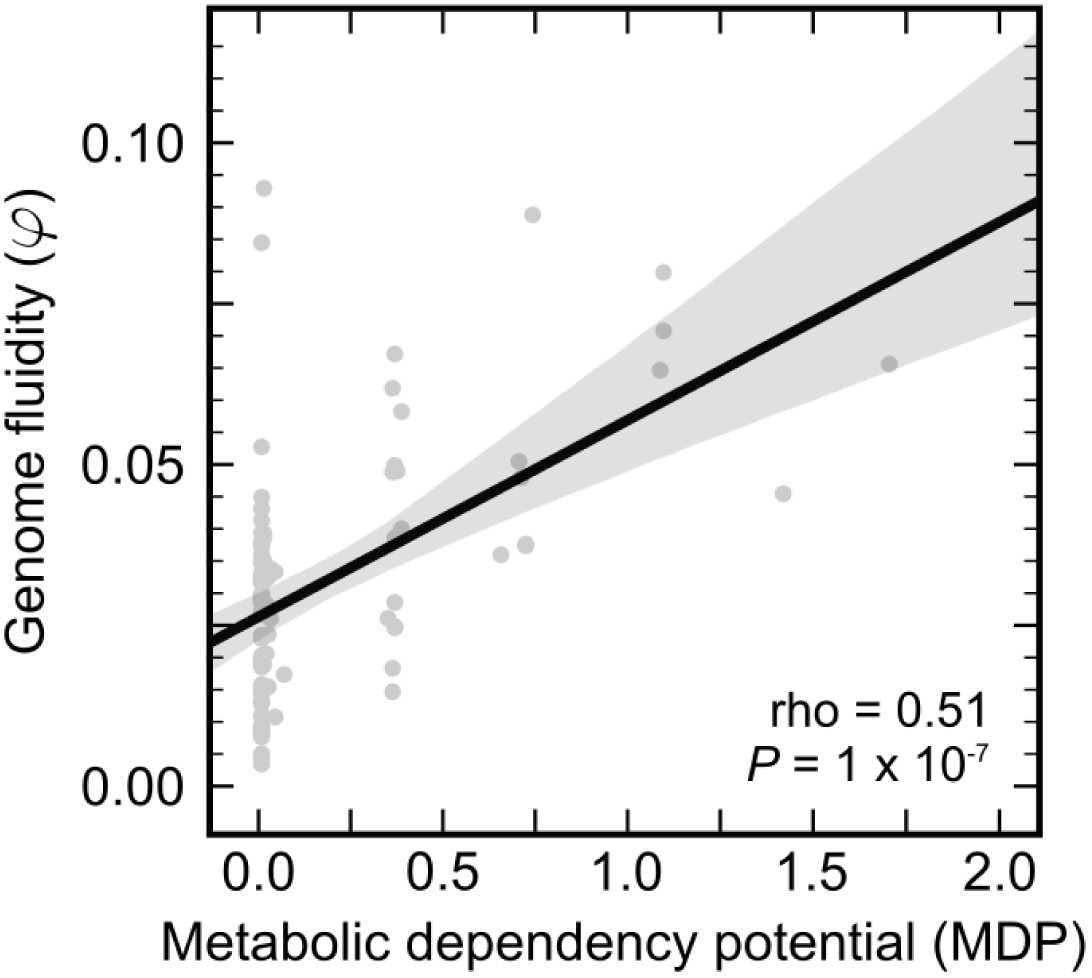
The impact of considering medians instead of means for MDP. Same as figure 2a, except here to calculate *α*, instead of using the mean number of dependencies per strain across conspecific pairs, I considered medians. The solid black line represents a linear regression and the gray envelope around it, the 95% prediction interval. rho corresponds to Spearman’s nonparametric correlation coefficient and the *P* value to a one-way asymptotic permutation test for positive correlation. This choice does not significantly impact my results for MDP.

**Supplementary Figure 6.**
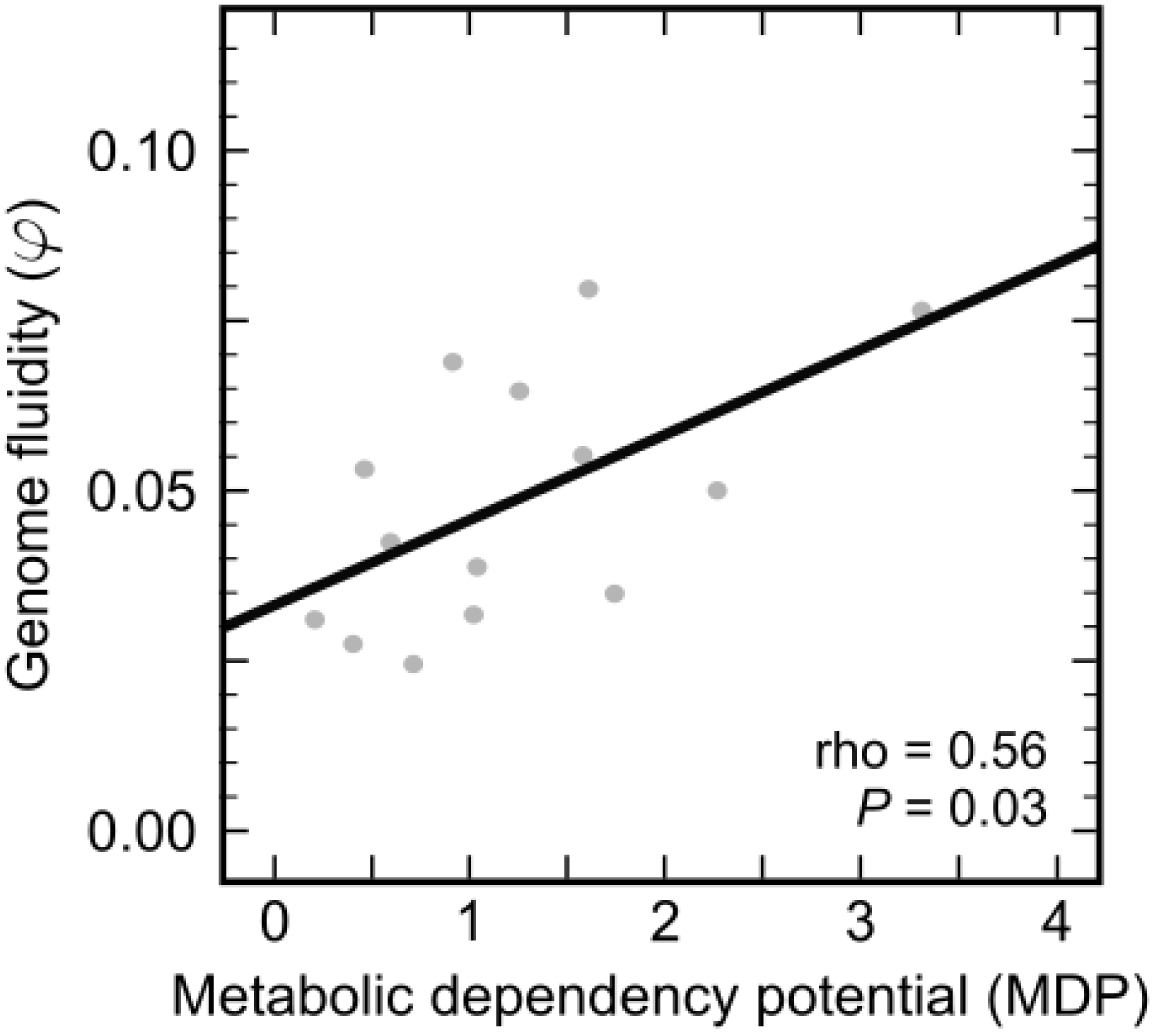
Co-occurring genomes from microbial community data capture my key observed trend. Scatter plot of genome fluidity *φ* versus conspecific metabolic dependency potential (MDP), similar to figure 2, but to test if the detected dependencies are realizable in nature, here I only included known co-occurring strains (see Methods). This resulted in 29 strains across 14 species (supplementary table 7). Each point represents the average number of dependencies detected per strain per condition across all co-occurring conspecific pairs for one species. The solid black line represents a linear regression and the gray envelope around it, the 95% prediction interval. MDP still increases significantly with increasing genome fluidity.

**Supplementary Figure 7.**
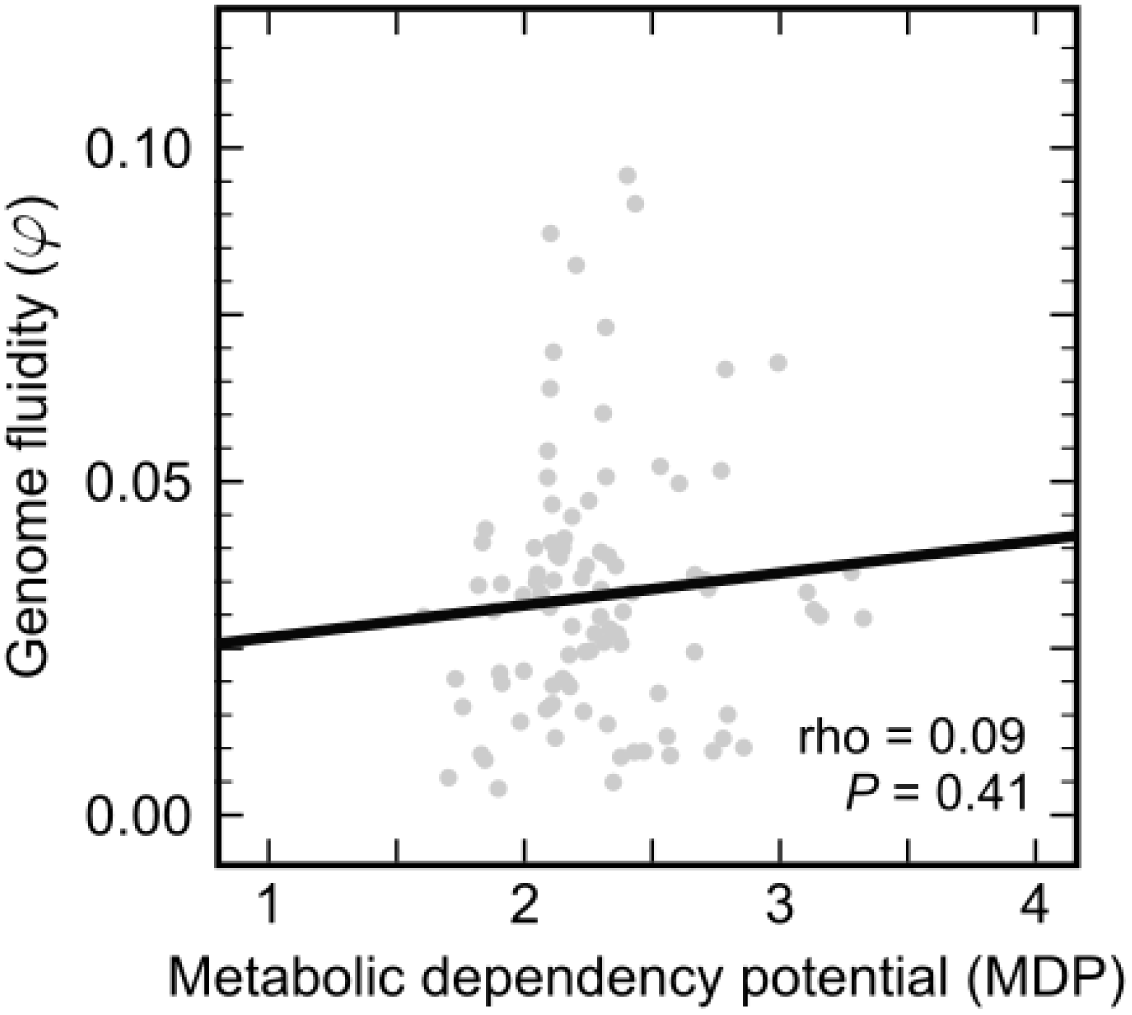
Complementary metabolic interactions are also likely to be common between prokaryotic species. Scatter plot of genome fluidity *φ* versus conspecific metabolic dependency potential (MDP), similar to figure 2, but here MDP was measured between inter-specific strains (see Methods). For each strain within a species, I measured its dependency potential with 25 randomly chosen strains from other species. Each point represents the average number of dependencies detected per strain per condition across several inter-specific pairs for one species. The solid black line represents a linear regression and the gray envelope around it, the 95% prediction interval. MDP is nonzero for all species in the study, though it does not increase significantly with increasing genome fluidity.

**Supplementary Figure 8.**
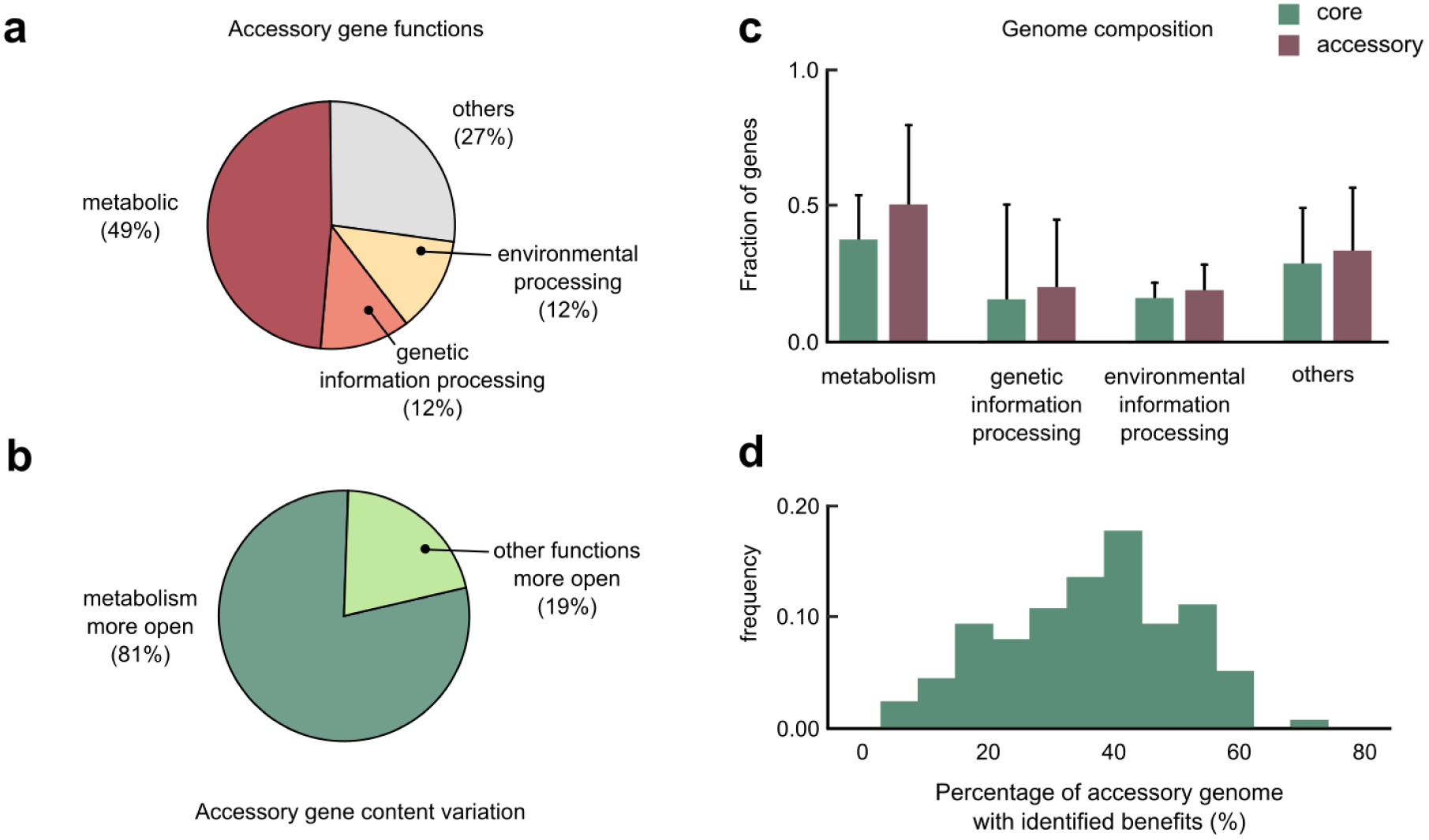
Quantitative aspects of prokaryotic core and accessory genomes. **a,** Pie chart of the average fraction of genes belonging to different functions typical to prokaryotic accessory genomes (for the 1,339 genomes in this study). **b,** Pie chart of the fraction of 96 species in the study where metabolic functions showed more gene content variation than other functions in accessory genomes (namely, genetic information processing, environmental information processing, and others). **c,** Bar chart of the typical functional composition of both core and accessory genomes, for the 96 species in the study. Values indicate median fraction, and error bars indicate the extent of variation observed across the genomes studied. Metabolic functions are enriched in accessory genomes when compared with core genomes. **d,** Histogram of the fraction of accessory genes in each strain, which I identified as potential contributors to metabolically beneficial functions in the study.

**Supplementary Figure 9.**
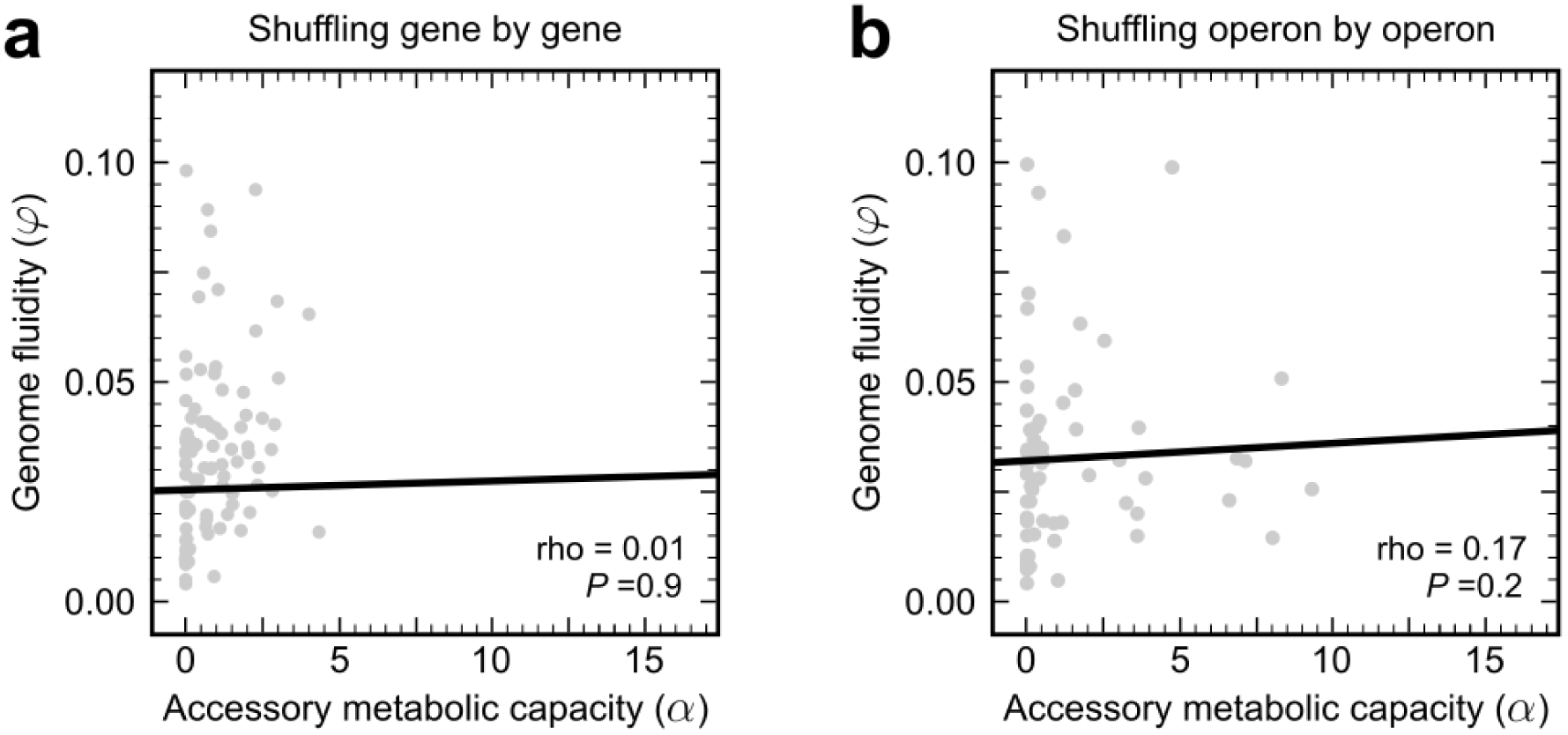
Randomly shuffling species pangenomes significantly diminishes *α*. Scatter plot of genome fluidity *φ* versus accessory metabolic capacity *α* for the prokaryotic species in this study, with each species’ accessory content randomly shuffled between strains, either **a,** gene by gene (for 96 species), or **b,** operon by operon (for 64 species, see Methods). The solid black lines represent linear regression. In both cases, not only does randomly shuffling accessory genes significantly reduce the additional biosynthetic potential of each strain’s accessory genome, there is no significant correlation with increasing accessory content (compared with figure 1).

**Supplementary Figure 10.**
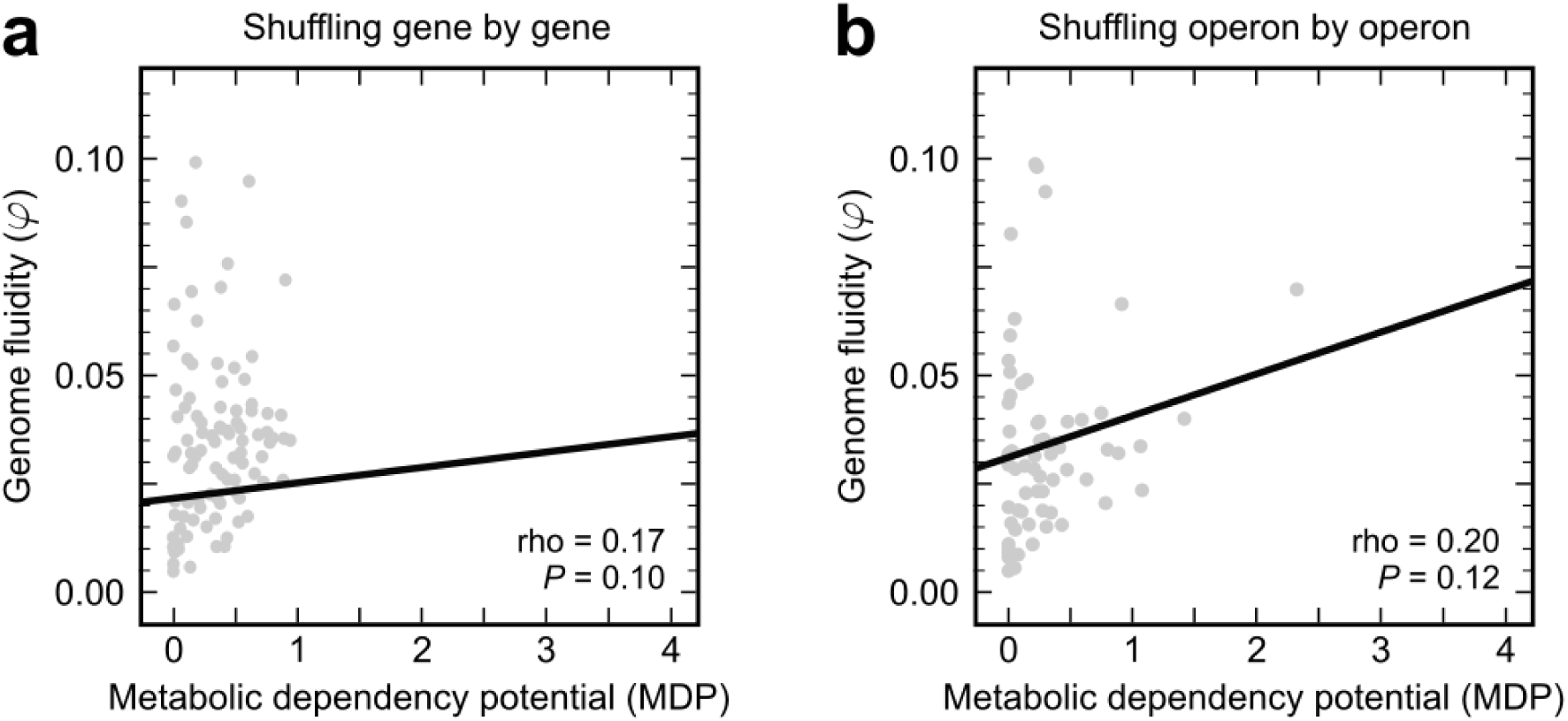
Randomly shuffling species pangenomes significantly diminishes MDP. Scatter plot of genome fluidity *φ* versus metabolic dependency potential MDP for the prokaryotic species in this study, with each species’ accessory content randomly shuffled between strains, either **a,** gene by gene (for 96 species), or **b,** operon by operon (for 64 species, see Methods). The solid black lines represent linear regression. In both cases, not only does randomly shuffling accessory genes significantly reduce the number of metabolic dependencies between conspecific pairs, there is no significant correlation with increasing accessory content (compared with figure 2a).

**Supplementary Figure 11.**
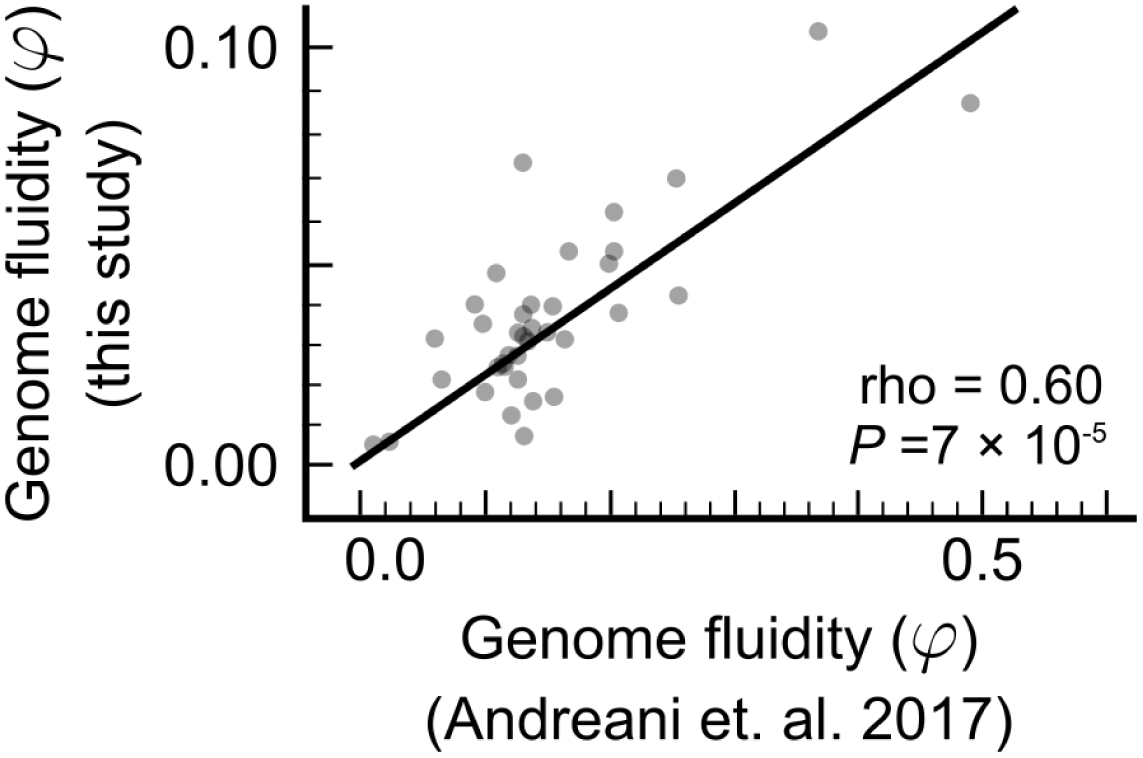
Comparing KEGG ortholog-based genome fluidity estimates with alignment-based estimates. Scatter plot of genome fluidity *φ* using KEGG (this study; y-axis) versus estimates of *φ* reported in Andreani et. al., *The ISME Journal* (2017) (x-axis) for the 38 species common to both analyses. Each point represents one species. My measure uses KEGG’s orthologous gene categories for scoring gene presence-absence, whereas Andreani et. al. use thresholded sequence alignment to classify genes into “families”. The solid black line represents a linear regression. rho corresponds to Spearman’s nonparametric correlation coefficient and *P* to a one-way asymptotic permutation test for positive correlation. While my estimates are lower than those of Andreani et. al., they show consistent linear scaling and can thus still be used to infer trends.

## Supplementary Tables

**Supplementary Table 1:** List of all 96 species and 1,339 strains used in the study

**Supplementary Table 2:** List of all 59 carbon sources used as nutrients

**Supplementary Table 3:** List of all 30 compounds assumed to present in all conditions

**Supplementary Table 4:** List of all 137 key biomass precursors used to define biosynthetic potential

**Supplementary Table 5:** Summary table with all measurements made during the analyses

**Supplementary Table 6:** List of all 562 Model SEED-derived gap-filled reactions added to the KEGG metabolic models

**Supplementary Table 7:** List of all 29 conspecific strains found to co-occur in microbial community data

**Supplementary Table 8:** List of all 795 strains used for operon shuffling

